# Higher order analysis of gene correlations by tensor decomposition

**DOI:** 10.1101/579276

**Authors:** Farzane Yahyanejad

**Affiliations:** Department of Computer Science, University of Illinois at chicago

**Keywords:** Tensor decomposition, Signal processing, Higher order correlation, Signal transduction, Yeast Saccharomyces

## Abstract

This study advances our understanding of inter- and intra-pathways higher order signaling in the cellular system and it leads to new discovery of multiple intracellular structures in signal transduction pathways in *yeast Saccharomyces*. We present a new tensor decomposition algorithm in reconstructing the pathways based on higher correlations among genes that compose a cellular system. The higher order gene correlation (HOGC) analysis has the power to elucidate gene’s higher interaction dependencies which has been barely understood. Recent studies i.e. [24] have experimentally revealed that multiple signaling proteins, yet sometimes infinite, may assemble to meaningful structure to transmit a receptor activation information. In this paper we reveal 3-order genomic correlations among significant component of the cellular system. This is the first time such a systematic and computational model provided for analysis of higher order correlations among genes. We use new fast algorithm to formulate a genes × genes × genes × decorrelated rank-1 sub-tensors (complexes) which can be associated with functionally independent pathways. Then we model higher order tensor decomposition 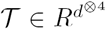 which is constructed by *K* tensors of genes × genes × genes. Each new tensor is constructed by an orthogonal projection of data signal onto a designated basis signal to keep common sub-tensors in both signals. Our model for decomposing tensor order-4 approximates series of tensors as linear components of deccorelated rank-1 sub-tensors over tensor of order-3 and rank-3 triplings among sub-tensors. The linear components represent intra-pathway in cell signaling and triplings implicate inter-pathways higher order signaling. Through structural studies of inter- and intra-higher order signaling pathways, we uncover different scenario that involves triple formation of signaling proteins into higher order signaling machines for transmission of receptor activation information to cellular responses.

## 1 Introduction and motivation

The biological functioning and life of a cellular system is controlled by signaling and energy transfer interactions among its numerous constituents such as proteins, RNAs, DNAs and other small molecules, and usually involve a *cascade* of biochemical reactions or other physical interactions among these constituents. Consequently, cellular systems generate genomic signals, such as mRNA expression and DNA-bound proteins’ occupancy levels, that can be measured experimentally using various methods such as DNA microarrays. Often, biologists model such a cellular system by presenting the interaction data in the form of a network diagram (some type of graph), *optionally* with some mathematical formulation of its dynamics. Usual mathematical formulations of the dynamics typically assume that each node *u* in the network has an associated “state” variable (representing, for example, concentration of the corresponding protein) that is a function of the “time” variable *t*, and describes how the *value* of this variable at a node (“state” of the node) depends on the state of the nodes interacting with it. A major drawback of using such graph-theoretic tools on a single network diagram lies in ignoring the time or ignoring the *higher-order* correlations of the interactions which may lead to *inaccurate* or *incomplete* analysis. For example, a network diagram only encodes pairwise correlations of node state variables, and thus cannot represent a joint *k*-way correlation among *k* state variables for any *k >* 2. If precise equations of time evolutions of state variables are given then we could of course completely ignore the network diagrams and work with the given equations, but then we lose the advantage of employing graph-theoretic tools and instead fall back on analysis techniques which are often *hard* to employ effectively because of difficulties of estimating precise equations and the non-trivial *non-linear* natures of these equations. The main topic of this paper is to study higher order (*beyond pairwise*) genomic correlations among significant components (*e.g.*, genes) of the cellular systems based on the measured signaling data. This is the first time such a mathematical model is given to uncover structural assemblies of higher order correlations among genes. Higher-order correlations among genes conditioned by the biological and experimental settings in which they are observed are *not* yet very well understood because of the difficulties to detect them genetic mapping studies, and not too many examples of such correlations have been described in the literature. However, evidences from model organisms suggest that these higher-order correlations among genes contribute *frequently* to genetic studies [9, 12, 15, 22, 26]. Informally, the main goal of this paper is to use powerful *tensor analysis methods* to provide the foundations of *systematic* and *computationally efficient* approaches to distinguish significant higher order correlations among the elements of biological systems.

For genes × genes networks, several previous researchers have proposed and analyzed models that separate genome-scale signals (*e.g.*, mRNA expression) into mathematically defined patterns that correlate with the independent biological and experimental processes and cellular states that compose the signals [1–3]. In this paper, we take this approach one step further to detect three-way correlations between the components of the biological system using tensor decompositions. These higher relations among the activities of genes may conditioned by the biological and experimental settings in which they are observed. We call them higher order signaling pathways. For example, the mRNA expression patterns of the yeast Sacchromyces genes *CIS*3, *SWI*4 and *HTA*1 have higher order correlations during cell-cycle progression. A single genomic signal matrix of correlations cannot describe differences in such higher level relations. More specifically, in this paper we describe the usage of higher order tensor decomposition via fast spectral algorithms (based on power iterations) to reconstruct pathway-dependent relations among the genes of a cellular system from higher-order correlations represented as symmetric tensors from measured genomic signals. Our method computes a genes × genes × genes tensor from signaling data, and decomposes the tensor to a *series* of genes × genes × genes decorrelated rank-1 sub-tensors. The rank-3 triplings among these sub-tensors are associated with higher-order inter-pathway correlations of the signaling pathways. To accurately approximate the higher-order correlations of the genes, we use a series of only those sub-tensors that are *common* to both signals by using the Moore-Penrose pseudo-inverse projection; this projects the signaling data onto a designated “basis” signal and simulate observations of only those pathways that are manifest under both sets of the biological and experimental conditions in which the data and basis signals are measured. Sub-tensors and tripling of sufficiently high significance represent independent pathways or higher-order correlations among them common to all or exclusive to a subset of the signals, and discretized sub-tensors and their triplings may highlight unknown differentials (*i.e.*, pathway-dependent relations) between genes. We illustrate our new fast decomposition algorithm and the new approach for finding higher-order correlations by analyzing the mRNA expression data of all common genes with manifested stages in the cell cycle of yeast (*S. cerevisiae*) [6] and DNA-binding data of the yeast transcription factors that are involved in cell-cycle, development, and biosynthesis programs [16] in pheromone-response classifications.

## 2 Method

Following widely used conventions, we will denote tensors, matrices and column vectors by calligraphic uppercase letters (*e.g., 𝒳*), boldface lowercase letters (*e.g.*, **x**) and non-bold lowercase letters with overhead arrows (*e.g.*,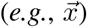), respectively. Individual elements will be denoted by the corresponding (non-bold) lower-case letter with appropriate indices, *e.g., x*_*i,j*_ for matrix **x**. We assume that the reader is familiar with the basic concepts and definitions associated with tensor analysis such as in [13, 14, 18].

### 2.1 Mathematical Models

Let the *symmetric* tensor 𝒯^(1)^ of size *N* × *N* × *N,* where *N* is the number of genes, tabulate the non-directional tensor of three-way correlations among the genes of a cellular system. We say tensor 𝒯^(1)^ is *symmetric* if the entries 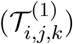 are invariant under permuting the indices. The tensor 𝒯^(1)^ is computed from a genome-scale signal, referred to as the data signal, of mRNA expression levels (or similar other data). Assume that these expression levels come from measurements conducted in a set of *M*_1_ samples of the cellular system and are subsequently tabulated as a *N* × *M*_1_ matrix **a**^(1)^, such that each entry of the tensor is:

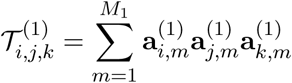

To find significant gene components of the correlation tensor, we decompose the tensor to *M*_1_ series of rank-1 sub-tensors of size *N* × *N* × *N* by a fast spectral algorithm (to be described later):

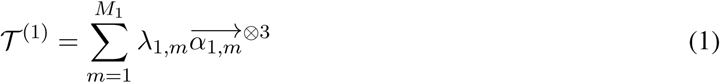

where *λ*_1,*m*_ is a scalar, ⊗denotes the tensor product operation, 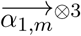 is a short-hand notation for 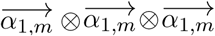 and the *m*^th^ sub-tensor 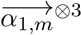 is the 3^rd^ order tensor product of the *m*^th^ gene component 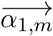. The *M*_1_ gene-values *λ*_1,*m*_’s and *M*_1_ gene-components 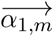 define the *M*_1_ non-negative higher-order gene level of gene expressions such that the expression of the *m*^th^ gene-value in the *m*^th^ gene-component is the *m*^th^ gene expression of subspace of 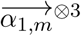. The significance of the *m*^*th*^ sub-tensor (*α*_1,*m*_ ⊗ *α*_1,*m*_ ⊗*α*_1,*m*_) is indicated by the *m*^*th*^ “fraction of gene-value”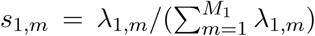 which is the higher expression correlation captured by *m*^*th*^ sub-tensor relative to that captured by all sub-tensors. Each sub-tensor is decorrelated of all other sub-tensors since by our algorithm we find orthogonal components under a small amount of error.

## 3 Results

### Tensor order 3 decomposition

In this part we give an algorithm that finds gene-components and gene-values in (1) with presence of very small spectral norm error. Our algorithm is inspired by orthogonal tensor decompositions proposed by Anandkumar et al. [4] and Ma et al. [11]. In general, we want to recover the gene-values *λ*_*k,m*_ and gene-components 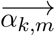 for *M*_*κ*_ samples of cellular system by finding a certain orthogonal tensor decomposition of every symmetric tensor 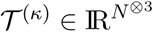 which is a 3^*rd*^ order tensor over IR^*N*^ such that 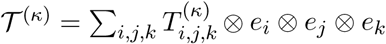 where *e*_1_, *e*_2_,…, *e*_*N*_ is the standard basis of IR^*N*^.

Decomposing general tensors is a delicate issue, tensors may not even have unique decomposition under a mild non-degeneracy condition. An orthogonal decomposition of a symmetric tensor 𝒯^(*κ*)^ is a series of orthonormal unit vectors 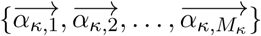 together with corresponding position scalars *λ*_*κ,m*_ ≥ 0 which in this paper we call them gene-value. For our algorithm we consider the case we have an orthogonally decomposable symmetric tensor 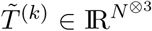 in tensor 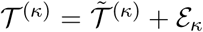, where *ε*_*k*_ is the perturbation in 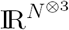 with small spectral norm. For a bipartition *P*_1_, *P*_2_ of the index set 𝒯^(*κ*)^, the spectral norm of the matrix unfolding ‖ 𝒯^(*κ*)^ ‖*P*_1,_*P*_2_ with rows indexed by the indices in *P*_1_ and columns indexed by indices in *P*_2_,

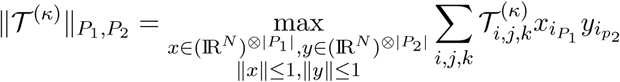

Since the tensor is *symmetric*, all possible spectral norms 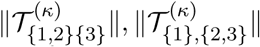, and 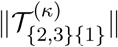 are the same. Also, since the order of our tensor is odd without loss of generality we can add the requirement that *λ*_*κ,m*_ is positive.

#### Algorithm 1 Fast spectral algorithm for 3^(*rd*)^ order tensor decomposition

1: **Input 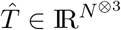**

2: Draw *U* ^(0)^ uniformly at random from IR^*N* ×*N*^

3: **for** *m* = 1 to log *N* **do**

4:

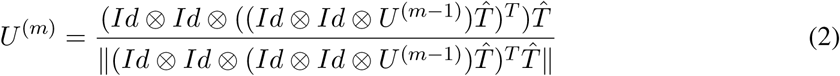

5: Initialize 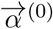 the top eigenvector of *U* ^(log*N*)^

6: **for** *m* = 1 to log *N* **do**

7:

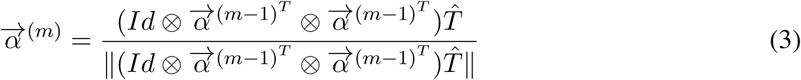

8: 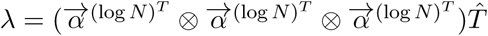

9: **return** the estimated eigenvector/eigenvalue pair 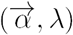

We remark that in our multilinear operation on tensors, we use the form 𝒯^(*κ*)^ ↦ (*U*_*i*_ ⊗*U*_*j*_ ⊗*U*_*k*_) 𝒯^(*κ*)^, where *U*_*i*_, *U*_*j*_, *U*_*k*_ are matrices with N columns and,

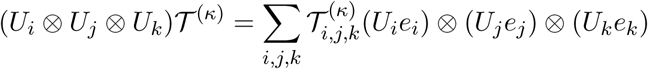

If some of the matrices of *U*_*m*_ are identity matrix, then the corresponding operation is called tensor contraction. For example, for 3^*rd*^-order tensor 𝒯^(*κ*)^ and matrix *U* ∈ IR^*N* ×*N*^, we call (*Id* ⊗*Id* ⊗*U)* 𝒯^(*κ*)^ the contraction of the third mode of 𝒯^(*κ*)^ with matrix *U* and the result of this contraction is a vector. For vector 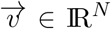, we call 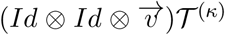 the contraction of the third mode of 𝒯^(*κ*)^ with vector 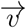 and the result of this contraction would be matrix.

The key step of the algorithm to find the single component of the tensor 𝒯^(*k*)^ is repeating two contractions of the tensor under power iteration in (2). We first generate matrix *U,* a uniform *N* ×*N* dimensional matrix and contract tensor 𝒯^(*κ*)^ with *U* to get a vector. Then we contract the tensor 𝒯^(*κ*)^ by contracted 𝒯^(*κ*)^ one more time and we get a matrix. Let us assume the result of the contraction of the tenor *T* ^(*κ*)^ with *U* is vector 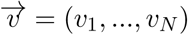,

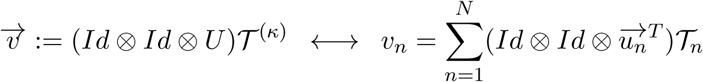

where 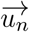 is *n*^*th*^ column of the matrix *U* and 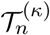 is *n*^*th*^ slice of tensor *T*^(*κ*)^. Now we contract tensor *T*^(*k*)^ by vector 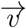 one more time by the following contraction and find matrix *U* which we normalize and use it in our power iteration update in (2),

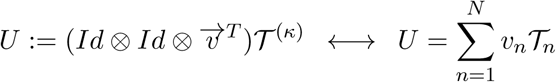

Since we have two contractions at each step and updated result would be a matrix instead of a vector in (2), Algorithm 1 establishes the convergence for extracting a single component of the orthogonal decomposition faster than the convergence of tensor power method for orthogonal decomposition in [4]. After log *N* power iteration we get converged matrix *U*. Then we run log *N* power iteration on top eigenvector of matrix *U* in (3) and the final output would be 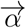, the single component of the tensor. The algorithm succeeds with probability at least 1/log *n* over the randomness of the algorithm when the ration between the largest and second largest eigenvalue of contracted tensor is at least 1 + 1*/* log *n*. We can find full analysis of similar method is in [11]. To find the corresponding *λ*_*κ,m*_ we contract tensor 𝒯^(*κ*)^ in all 3 ways by the component. By running Algorithm 1 *M*_*κ*_ times we can extract an approximated decomposition of 𝒯^(*κ*)^.

Computation of subsequent gene-component can be computed with deflation 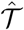, i.e., 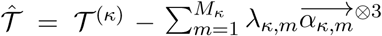.

### Tensor order 4 decomposition

Let the fourth-order tensor 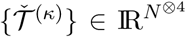 of size *K* × *N*-genes × *N*-genes × *N*-genes tabulates a series of *K* genome scale order-3 tensors 𝒯^(*κ*)^,

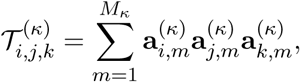

such that each tensor is constructed by a new genomic signal (i.e. DNA-bound protein’s occupancy levels) which is created by Moore-Penrose pseudo-inverse projection of genome scale signal **a**^(1)^ on designated basis signal **p**^(*κ*)^ of size *N*-genes × *M*_*κ*_-arrays, of, e.g., proteins’ DNA binding levels measured in a set of *M*_*κ*_ samples of the cellular system,

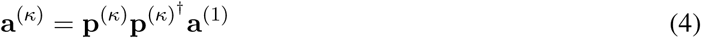

By this computation we want to keep only those sub-tensors which are common to both signals **a**^(1)^ and new projected signal **a**^(*κ*)^. Now we define and compute a higher order tensor decomposition of the tensor of tensors 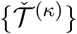,

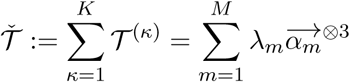

The result of above decomposition are gene-components 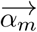 and gene-values *λ*_*m*_which are decorrelated in overall tensor 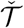 as our algorithm find the orthogonal components with good approximation. These deccorelated gene-components in overall tensor might not be decorrelated in any one of individual tensors, since *λ*_*κ,m,l,n*_ ≠ 0. Here we propose a new model based on higher order tensor decomposition above which formulates each individual tensor order-3 𝒯^(*κ*)^ as a series of 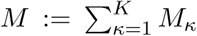 rank-1 symmetric decorrelated sub-tensors and the series of 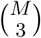 rank-3 tripling among these sub-tensors such that,

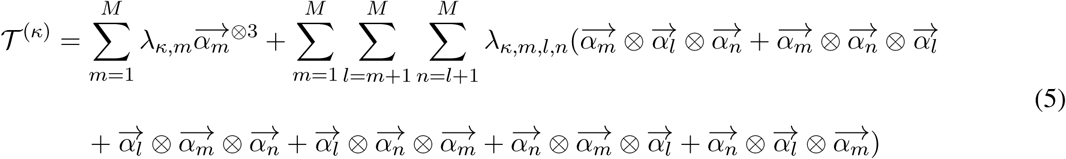

for all *κ* = 1, 2, *…K*. Tripling of components 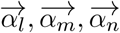 is sum of tensor product of all combinations of these sub-tensors for all *m* ≠ *l* ≠ *n*. By this definition we can find the significance of HOGC *m*^*th*^ sub-tensor in the *κ*^*th*^ tensor. The significance is indicated by the *m*^*th*^ fraction of gene-component of the *κ*^*th*^ tensor 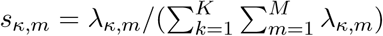, i.e., the correlation of components captured by the *m*^*th*^ sub-tensor in the *κ*^*th*^ tensor relative to that captured by all sub-tensors. Similarly, the amplitude of the fraction 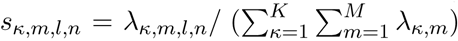 indicates the significance of the tripling among the *m*^*th*^, *l*^*th*^ and *n*^*th*^ sub-tensors in the *κ*^*th*^ tensor.

### 3.1 Biological interpretation of Yeast intracellular signaling pathways in higher order assemblies

To determine HOGC inter- and intra-pathways in Yeast cellular system, we compute *K* = 4 HOGC tensors by signal matrices **a**^(1)^, **a**^(2)^, **a**^(3)^, and **a**^(4)^. The genomic signal matrices **a**^(2)^, **a**^(3)^, and **a**^(4)^ are driven by projection **p**^(1)^, **p**^(2)^ and **p**^(3)^ onto basis signals **a**^(1)^ (4), which tabulates the relative binding DNA-bound protein occupancy levels of the yeast transcription factors that are involved in cell-cycle, development, and biosynthesis programs respectively under experimental conditions of Spellman et al. and Roberts et al. [16, 20, 21]. We compute all tensors 𝒯^(1)^, 𝒯^(2)^, 𝒯^(3)^, and 𝒯^(4)^ on common genes among **a**^(1)^, **p**^(1)^, **p**^(2)^ and **p**^(3)^ with manifested stages for either pheromone or cell cycle classification which tabulates relative mRNA expression levels of 27 yeast genes with valid data in the number of presented samples of a cell cycle time course of a culture synchronized by the mating pheromone *α* factor. Before computing 𝒯^(1)^ from the data signal **a**^(1)^, we use IterativeSVD to fill missing data in **a**^(1)^ and to approximately center the expression pattern of each gene in it’s invariant level (mean). IterativeSVD is matrix completion by iteratively low-ran svd decomposition which is similar to SVDimpute from [Missing value estimation methods for DNA microarrays] [23, 27]. Tensors 𝒯^(2)^, 𝒯^(3)^, and 𝒯^(4)^ are computed by **a**^(2)^, **a**^(3)^, and **a**^(4)^ respectively.

For both pheromone and cycle classifications we associate sub-tensors and the tripling among them with most likely expression correlations based on the *P*-values calculated by hypergeometric probability distribution of the *Q* triples of annotations among *T* which is total number of triples,

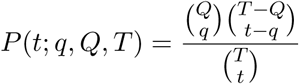

By combinatorics and assuming a hypergeometric probability distribution [19], we can find the probability of a specific annotation in *Q* triples of genes’ annotations from a population of *N* genes that contains exactly *T* triples with that annotation, where in each annotation is one of the possible combination of triple among cell cycle and pheromone response states. The association of all sub-tensors of projected tensors to annotations from two different classifications of cell-cycle and pheromone-response are brought in Table1.

**Table 1:**
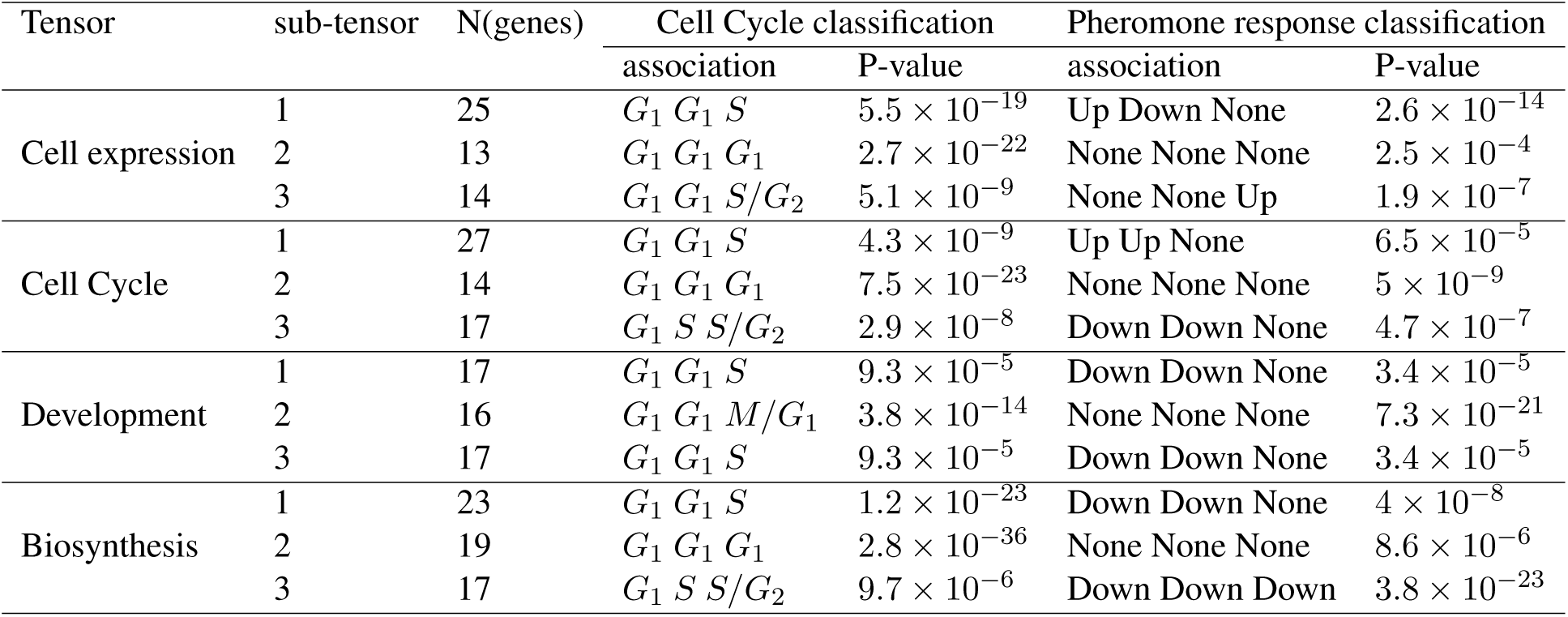
Most likely parallel associations of the most three significant independent higher order of signaling pathways and number of participated genes in each pathway of tensor 𝒯^(1)^ created by data signal a^(1)^, 𝒯^(2)^ created by a^(2)^ the projection of a^(1)^ onto the cell cycle, 𝒯^(3)^ created by a^(3)^ the projection of a^(1)^ onto the development, and 𝒯^(4)^ created by a^(4)^ the projection of a^(1)^ onto the biosynthesis basis signals, according to the traditional and microarray classifications of cell cycle- and pheromone-regulated yeast genes.

| Tensor | sub-tensor | N(genes) | Cell Cycle classification |  | Pheromone response classification |  |
| --- | --- | --- | --- | --- | --- | --- |
|  |  |  | association | P-value | association | P-value |
| Cell expression | 1 | 25 | $G_1 G_1 S$ | $5.5 \times 10^{-19}$ | Up Down None | $2.6 \times 10^{-14}$ |
| | 2 | 13 | $G_1 G_1 G_1$ | $2.7 \times 10^{-22}$ | None None None | $2.5 \times 10^{-4}$ |
| | 3 | 14 | $G_1 G_1 S/G_2$ | $5.1 \times 10^{-9}$ | None None Up | $1.9 \times 10^{-7}$ |
| Cell Cycle | 1 | 27 | $G_1 G_1 S$ | $4.3 \times 10^{-9}$ | Up Up None | $6.5 \times 10^{-5}$ |
| | 2 | 14 | $G_1 G_1 G_1$ | $7.5 \times 10^{-23}$ | None None None | $5 \times 10^{-9}$ |
| | 3 | 17 | $G_1 S S/G_2$ | $2.9 \times 10^{-8}$ | Down Down None | $4.7 \times 10^{-7}$ |
| Development | 1 | 17 | $G_1 G_1 S$ | $9.3 \times 10^{-5}$ | Down Down None | $3.4 \times 10^{-5}$ |
| | 2 | 16 | $G_1 G_1 M/G_1$ | $3.8 \times 10^{-14}$ | None None None | $7.3 \times 10^{-21}$ |
| | 3 | 17 | $G_1 G_1 S$ | $9.3 \times 10^{-5}$ | Down Down None | $3.4 \times 10^{-5}$ |
| Biosynthesis | 1 | 23 | $G_1 G_1 S$ | $1.2 \times 10^{-23}$ | Down Down None | $4 \times 10^{-8}$ |
| | 2 | 19 | $G_1 G_1 G_1$ | $2.8 \times 10^{-36}$ | None None None | $8.6 \times 10^{-6}$ |
| | 3 | 17 | $G_1 S S/G_2$ | $9.7 \times 10^{-6}$ | Down Down Down | $3.8 \times 10^{-23}$ |

The Cell cycle classification includes five cellular states (*G*_1_, *S, G*_2_, and *M)* [8]. For pheromone response classification we have either state Up or Down [10]. The stages might be None which means we don’t know about the state of that gene regarding the classification. We select *t* = 120 higher correlations among *T* = 2925 triples of *N* = 27 genes at the intersection of basis signals to find the association of most likely cellular states with each sub-tensor and tripling among them for both classifications.

#### Significant independent sub-tensors associated with intra-pathways in higher order signaling machines

In this part we analyze genes are pathway dependent in higher order signaling machines. Genes are involved in independent intra-pathway higher order signaling are assembled to the structure regarding of their cell-cycle phases. The algorithm also drives the impact of each independent higher order signaling pathway in overall multiple intracellular signaling. The most three significant sub-tensors which our algorithm uncovers on 𝒯^(1)^, capture ≈ 67%, 5%, and7% respectively of the higher expression correlation of 𝒯^(1)^. There are independent pathways associated to the HOGC sub-tensors which are manifest in the data signal **a**^(1)^. The result follows by he *P* values for the distribution of the 10 genes. The genes microarray-classified as pheromone regulated and 25 genes which traditionally or microarray classified as cell cycle regulated genes with the subset of t=120 triples of genes with highest levels of expression correlation among all *T* = 2925 triples. We visualize these three discretized sub-tensors of the 𝒯^(1)^ that constitute 120 higher correlations in each sub-tensors with large amplitude among all high correlations Figure1. The result uncovers genes *MKC*7, *Y NK*1, *Y EH*1 and *GAS*2 are not pathway dependent in higher order signaling as they have not appeared in any HOGC sub-tensors.

**Figure 1:**
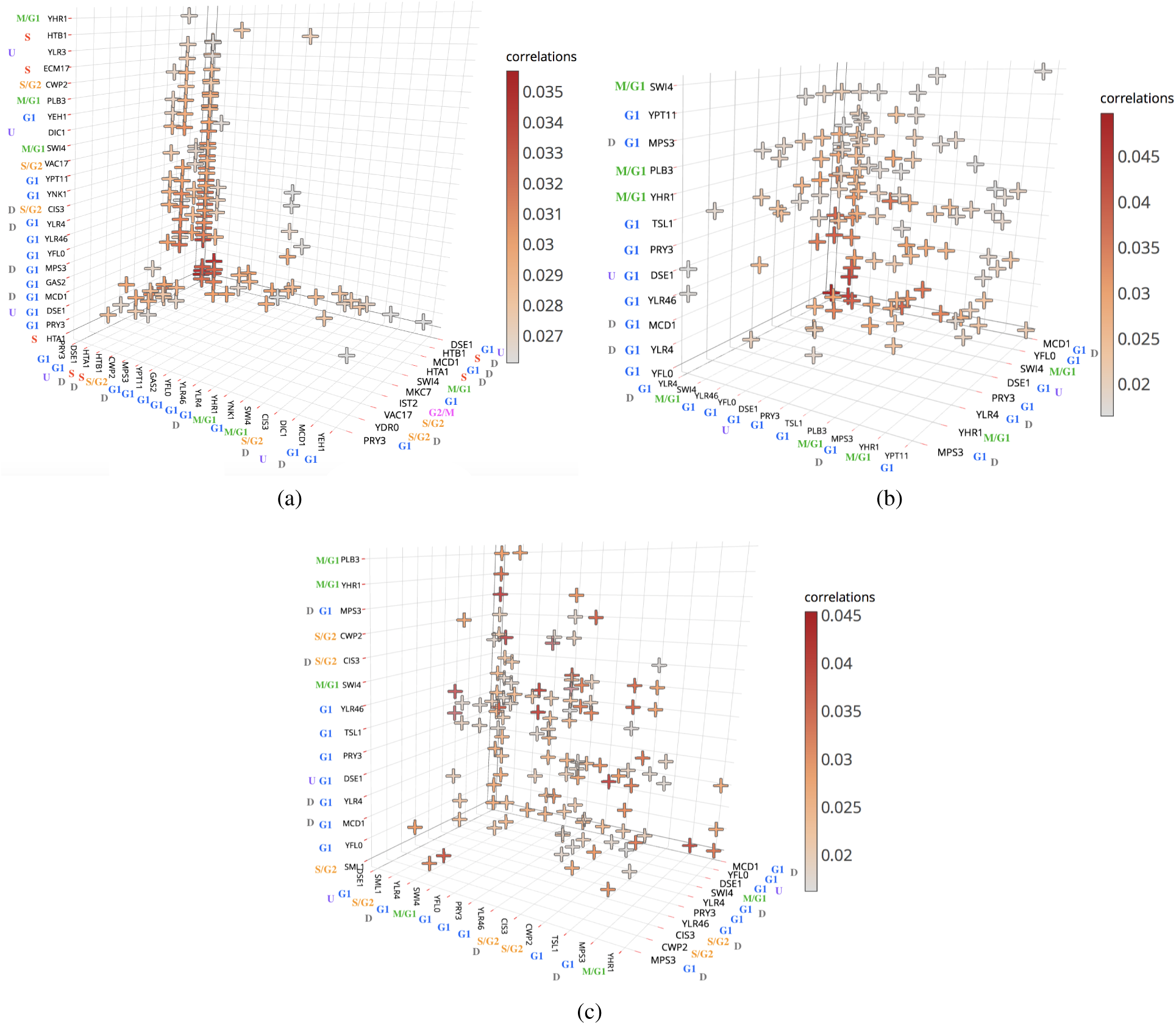
Discretized significant sub-tensors of the tensor 𝒯^(1)^ in the subset of 120 higher correlations largest in amplitude among all genes with their cell-cycle classificartions, *M/G*_1_, *G*1, *S, S/G*_2_, and *G*_2_*/M* and pheromone-response classifications, up-regulated and down-regulated. Most of the participated genes in higher order correlation among 27 genes in the intersection of DNA binding data are not reported to be regulated by pheromone but new structures of cell-cycle dependent among proteins can be observed. U and D are the gene code according to pheromone response classifications up-regulated and down-regulated respectively

Figure1a shows that the higher correlations among the genes in the first and most significant sub-tensor are pheromone-response independent, however we can see that all higher correlations, have at least one gene with cell-cycle stage *S* or *G*_1_. For example, *CIS*3, *SWI*4, and *Y HR*1 are not correlated since none of them peaks at either cell cycle *S* or *G*_1_, but we have HOGC among *CIS*3, *SWI*4, and *HTA*1 where *HTA*1 encodes cell cycle *S*.

In the second sub-tensor, The higher relations among the genes depend only on cell cycle classification phases *M/G*_1_ and *G*_1_ Figure1b. This sub-tensor discovers there is no higher correlation among genes with other classification phases, and therefore it filters out all decorrelated genes *IST* 2, *V AC*17, *Y DR*0, *HTB*1, *HTA*1, *CIS*3, *ECM* 17, *CWP* 2, *Y LR*3, *DIC*1, *SML*1, and *PLB*3 of 𝒯^(1)1^

The third HOGC sub-tensor omits those genes from cell cycle classification of *S* and *G*_2_*/M* since they are decorrelated from each other in higher order relations in this sub-tensor Figure1c. More precisely the higher relation of genes in this sub-tensor depends only on genes with cell cycle progression *G*_1_, *M/G*_1_, and *S/GG*_2_. For example, *CIS*3, *SWI*4, and *HTA*1 are not correlated since *HTA*1 encodes cell cycle *S*. Besides, genes presented in HOGC during mating are cycle dependent. Genes that are down regulated in response to pheromone are involved in the third higher order signaling intra-pathway if their cell-cycle stages are either *G*1 or *S/G*2. Also, We can see gene *DSE*1 with down regulated response to pheromone during mating is involved in all higher order correlations in signaling pathways. For all tensors, most likely parallel associations of the first three significant independent HOGC of signaling pathways and number of participated genes in each one are presented in Table 1.

**Table 2:** Most likely parallel associations of the sub-tensors of tensor 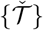 constructed by four tensors 𝒯^(1)^ 𝒯^(2)^ 𝒯^(3)^ and 𝒯^(4)^ and their tripling according to the traditional and microarray classifications of cell cycle- and pheromone-regulated yeast genes.

| sub-tensors or Triples between sub-tensors | Cell Cycle classification |  | Pheromone response classification |  |
| --- | --- | --- | --- | --- |
|  | association | P-value | association | P-value |
| 1 | $G_1 G_1 S$ | $8.5 \times 10^{-21}$ | Down Down Up | $7.4 \times 10^{-7}$ |
| 2 | $G_1 G_1 G_1$ | $1.2 \times 10^{-17}$ | None None None | $1.2 \times 10^{-4}$ |
| 3 | $S/G_2 S/G_2 S$ | $1.6 \times 10^{-18}$ | Down Down None | $4.8 \times 10^{-8}$ |
| 4 | $S/G_2 S/G_2 S$ | $1.6 \times 10^{-18}$ | Down Down None | $4.8 \times 10^{-8}$ |
| $1 \leftrightarrow 2 \leftrightarrow 3$ | $G_1 G_1 S$ | $5.6 \times 10^{-32}$ | None None None | $2.6 \times 10^{-15}$ |
| $1 \leftrightarrow 2 \leftrightarrow 4$ | $G_1 G_1 M/G_1$ | $9.6 \times 10^{-7}$ | None None Down | $5.1 \times 10^{-3}$ |
| $1 \leftrightarrow 3 \leftrightarrow 4$ | $S/G_2 G_1 S$ | $1.4 \times 10^{-13}$ | None None Down | $5.1 \times 10^{-3}$ |
| $2 \leftrightarrow 3 \leftrightarrow 4$ | $G_1 G_1 S$ | $4.4 \times 10^{-11}$ | None None Down | $1.6 \times 10^{-10}$ |

### sub-tensors and their tripling are associated with intra- and inter-pathways’ higher order signaling

Our model which is presented in (5) for higher order correlations among tensors 𝒯^(1)^, 𝒯^(2)^, 𝒯^(3)^, and 𝒯^(4)^ uncovers 4 significant sub-tensors and tripling among these sub-tensors which capture higher order correlations in each individual tensor Figure3. The significant decorrelated sub-tensors that are manifested in overall tensor capture ≈60%, 4%, and 6%, and ≪1% of the higher correlations among genes respectively. The associations of aforementioned sub-tensors and triplings among them is computed for 27 common genes (genes with manifested stage either in pheromone classification or cell-cycle) in **a**^(1)^, **a**^(2)^, **a**^(3)^, and **a**^(4)^ Table2.

The 3D visualization of four significant sub-tensors of higher order decomposition of individual tensors Figure2 reveals new hidden subsets of higher order correlation of intra-pathway genes. The sub-tensors are associated with the higher order signaling in independent pathways that are manifest in the overall tensor of higher correlations as well as the individual tensors Figure2. The higher order intra-pathway relation of genes in the first sub-tensor of overall decomposition brought in Figure2a.

**Figure 2:**
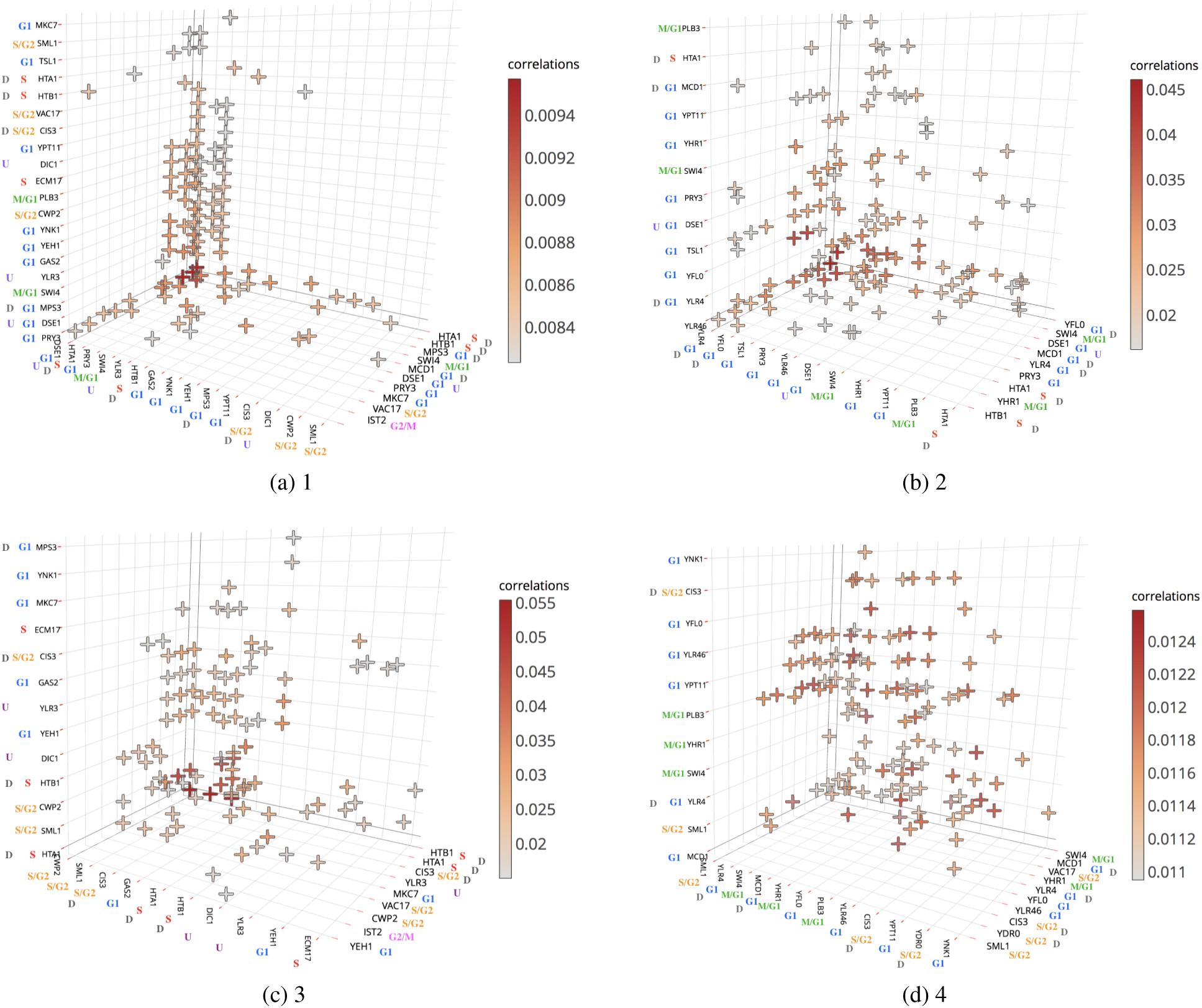
Discretized significant sub-tensors of the Tensor 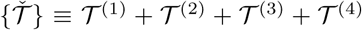 in the subset of 120 higher correlations largest in amplitude.

Figure2b uncovers that only genes with the cell cycle classifications of *G*_1_, *M/G*1, and *S* belong to this independent pathway in higher correlation. Genes with down regulated in response to pheromone that are involved in this higher order signaling intra-pathway depend on either cell-cycle phase *G*1 or *S*.

In the third sub-tensor, The higher relations among the genes depend only on cell cycle classification of *G*_1_, *S/G*_2_ and *S* Figure2c. Genes that are down regulated in response to pheromone are involved in the third higher order signaling intra-pathway if their cell-cycle stages are either *S* or *S/G*2.

The fourth sub-tensor keep those genes from cell cycle classification of *G*_1_, *S/G*2 and *M/G*1 Figure2d. This sub-tensor is also is pheromone dependent in the way that it highlights no higher correlation among involved genes are Up-regulated in pheromone-response classification. Genes that are down regulated in response to pheromone are involved in the third higher order signaling intra-pathway if their cell-cycle stages are either *G*1 or *S/G*2.

Figure3c,Figure3d shows the the contribution of the first sub-tensor to the higher expression correlation of development tensor 𝒯^(3)^ and biosynthesis tensor 𝒯^(4)^ which are constructed based on projected signals **a**^(3)^ and **a**^(4)^ is negligible, but it contributes to the higher expression correlation of tensor 𝒯^(1)^ and cell cycle tensor 𝒯^(2)^ Figure3a,3b. The second sub-tensor, which is associated with signal transduction pathway that only involves genes with cell cycle stage *G*_1_, *M/G*_1_, and *S* in higher order correlations contributes to the expression correlation of the higher correlations of tensor 𝒯^(1)^ and biosynthesis tensor 𝒯^(4)^ mostly Figure3a,Figure3d, and third sub-tensor to the higher expression correlation of tensor 𝒯^(1)^ and development tensor 𝒯^(3)^Figure3a,Figure3c. The most contribution of fourth sub-tensor which is associated with signal transduction pathway in higher order formations of genes with cell cycle *G*_1_, *S/G*_2_, and *M/G*_1_ is in cell cycle tensor 𝒯^(2)^ and development tensor 𝒯^(3)^ Figure3b,Figure3c, while it’s contribution to tensor 𝒯^(1)^ and biosynthesis tensor 𝒯^(4)^ is negligible Figure3a,Figure3d.

**Figure 3:**
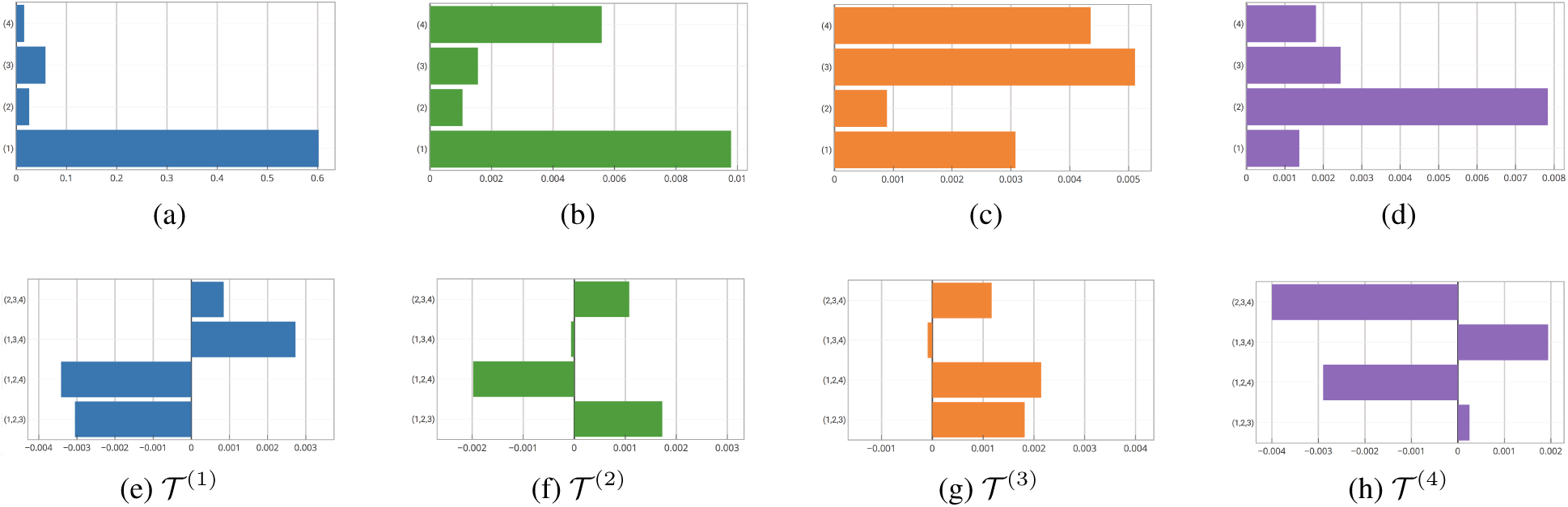
Significance of the sub-tensors of higher order tensor 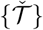 in each individual tensor 𝒯^(1)^, 𝒯^(2)^, 𝒯^(3)^, and 𝒯^(4)^ and the contribution of tripling among them.

Trough structural studies of inter-pathways signaling sub-tensors, we began to see different scenarios regarding formation of higher order signaling machines. For example, we identified this scenario that histones and putative proteins are involved in all higher order signaling of inter-pathways. The participated Histones in these HOGC inter-pathways’ higher order signaling are *HTA*1 and *HTB*1 and putative proteins are *Y LF* 0 and *Y LR*4 (*Y FL*0 and *Y LR*4 are short form of YORFs *Y FL*064*C* and *Y LR*462*W* since their gene’s name are unknown). The discretized sub-tensors and triplings highlight unknown higher order relations among genes that are in agreement with current understanding of the cellular system of yeast Figure4,Figure5. The tripling is associated with the higher order signaling among these independent pathways that are manifest in individual tensors.

**Figure 4:**
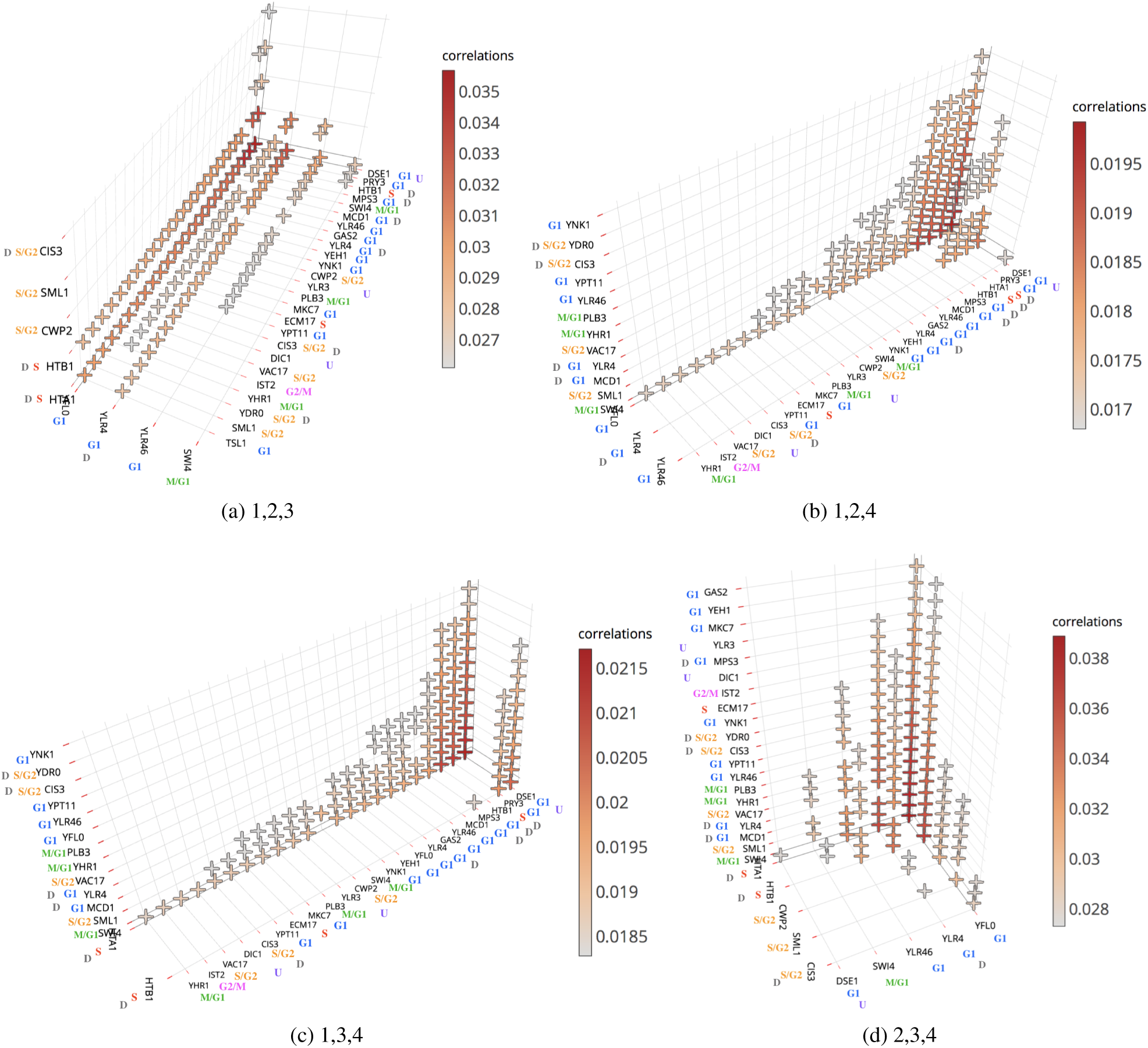
Discretized significant Tripling among the significant sub-tensors of Tensor 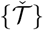 in the subset of 120 higher correlations largest in amplitude.

**Figure 5:**
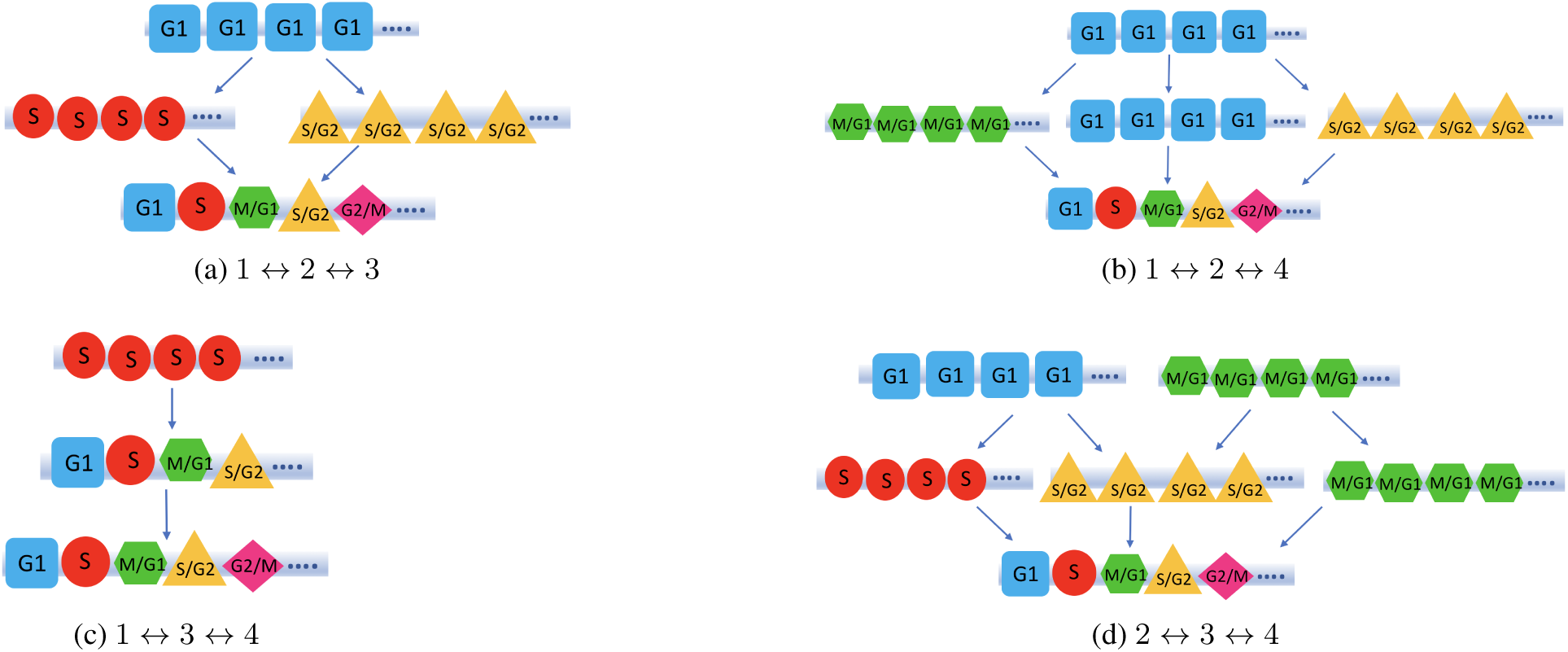
Higher order assemblies of cell cycle classification of genes in triplings among the significant HOGC sub-tensors of signal transduction Tensor 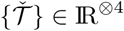 in cellular system in the subset of 120 higher correlations largest in amplitude. Each layer shows the stage of one of the involved genes in triple correlation for each tripling. For example, (a) shows that in first tripling among 𝒯^(1)^,𝒯^(2)^, and 𝒯^(3)^ the first gene is with cell-cycle stage *G*1, the second gene has cell-cycle stage *S* or *S/G*2, and the third one can be any of them.

The tripling among first, second and third sub-tensors is associated with *G*_1_ and one of the *S/G*_2_ or *S* and it has contribution in all individual tensors except 𝒯^(4)^ Figure3. To obtain comprehensive view of the relationship between higher order gene organization and induction of developmentally regulated gene-expression profile, our comparative analysis of intra-pathway of first, second and third reveals distinct set of cell cycle classification of (*G*_1_, *S*) or (*G*_1_, *S/G*_2_) in every triple of genes in this higher order level Figure5a. We might conclude that in higher correlation among tensor of expression correlations of 𝒯^(1)^, and development projected tensor of 𝒯^(3)^ gene with type *G*_1_ from cycle classification is an activation platform for correlation of other genes in this tripling. The most participated genes in this tripling is *Y FL*0, *Y LR,* and *Y LR*6 Figure4b.

The tripling among first, second and fourth sub-tensors reveals *G*_1_ as a premanent cell cycle association and it contributes in all individual tensors Figure3 Figure5b.The most participated genes in this tripling is *Y FL*0, *Y LR,* and *Y LR*6 Figure4a.

In all triples among first, third and fourth sub-tensors we have *S* and this tripling contributes in 𝒯^(1)^ and 𝒯^(4)^ while it’s contribution in individual tensors 𝒯^(2)^ and 𝒯^(3)^ is negligible Figure3g Figure3h. Figure4c shows the most significant genes in this tripling are *HTA*1, and *HTB*1. Therefore, this tripling highlights higher order correlations among the histones and other genes that are involved in triple of first, third and fourth sub-tensors.

The tripling among second, third and fourth sub-tensors which has contribution in 𝒯^(2)^, 𝒯^(3)^, and 𝒯^(4)^ reveals distinct subsets of *G*_1_, *S, G*_1_, *S/G*_2_, *M/G*_1_, *S/G*_2_ and *M/G*_1_, *M/G*_1_ Figure5d. Figure4d also shows the most significant genes in this tripling are *Y FL*0, *Y LR, HTA*1, and *HTB*1.

## 4 Discusion

Many questions are raised by higher order correlation analysis of genes, which need to be explored both theoretically and experimentally. Elucidating the biophysical principle governing the higher order signaling analysis among genes, regulated fashion may reveal the structural basis of clustering and uncover paradigm in cell singling. Higher-order assemblies may be an important aspect of many biological processes because they enable formation of precisely organized molecular machines from constituents present in inactive states at low concentrations to promote biochemical reactions in cells. We have defined tensors based on genome scale signal of, e.g., mRNA expression and proteins’ DNA-binding relations among genes and described a algorithm for efficient new tensor decomposition based on power iteration to separate genome scale nondirectional tensors into mathematically defined decorrelated components. We have demonstrated its usefulness on a three dimensional and four dimensional higher correlation data to uncover HOGC sub-tensors and HOGC tripling relations among them with clear biological and statistical significance. This approach tends to focus mainly on identifying higher order signaling in intra-pathways in cellular system and higher order signaling among these transduction signaling pathways (inter-pathway signaling) and also multiple intracellular structures of cell cycle genes. The proposed tensor decomposition was a simply approach to extract the orthogonal decomposition with high accuracy in analyses of higher order gene correlations in genome scale yeast data. These components and tripling among them include reconstruction of higher order signaling of inter- and intra-pathways from nondirectional tensor of higher correlations. The result uncovers new higher order coordinated differential relations mostly among cell-cycle and also some pheromone regulated genes in higher order pathway-dependent signalings of these genes. There are several interesting ways in which this model can be extended or changed. The model can be extended to higher-dimensional data or this factorization can be benefited multiple-tissue gene expression data sets.

## A Apendix

**Figure 6:**
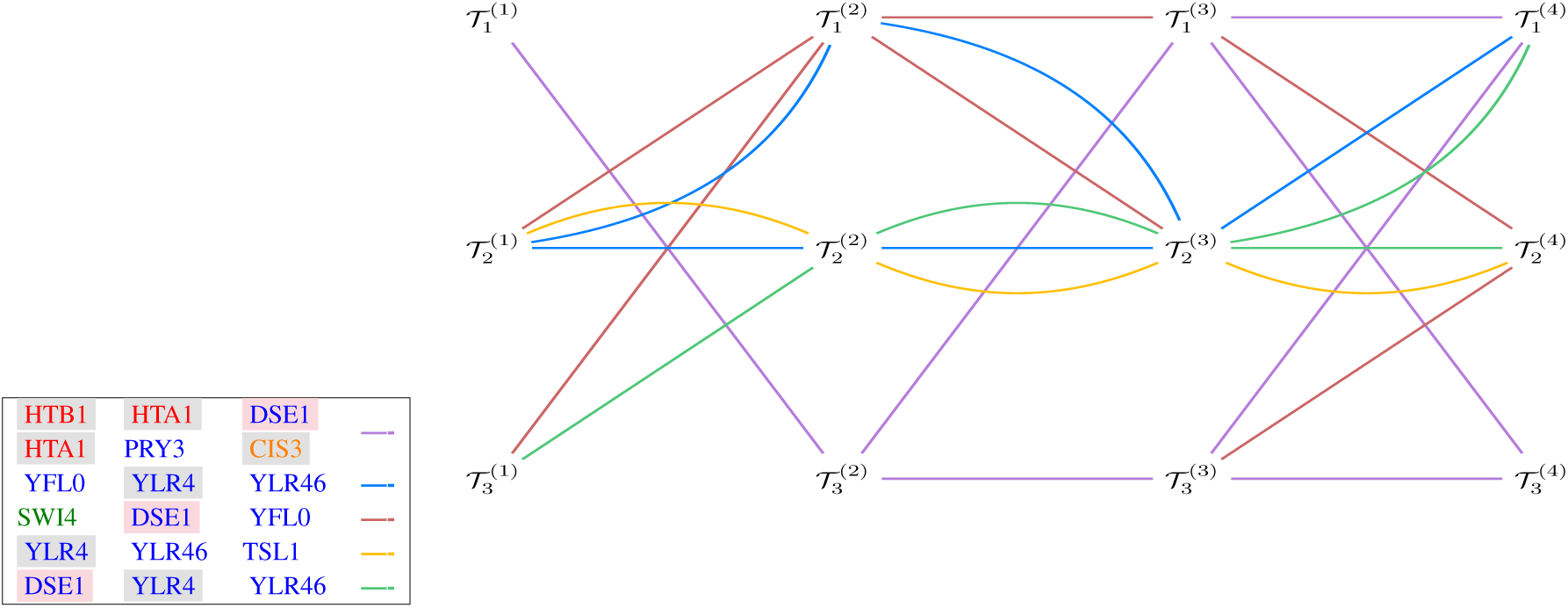
Common higher correlated genes among all sub-tensors of tensor 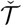. These higher order correlation of genes are not pathway dependent in higher order signaling machine. Color-coded for genes inside the box is according to their cell-cycle classifications, *M/G*_1_(green), *G*_1_(blue), *S*(red), *S/G*_2_(orange), and *G*_2_*/M* (pink), and separately according to their pheromone-response classifications, we highlight the up-regulated purple and down-regulated gray. The graph nodes are depicted in columns and rows. Each column in the graph shows HOGC sub-tensors of an individual tensor. For example 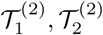,and 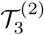 are most significant HOGC sub-tensors of tensor 𝒯^(2)^ which is Cell Cycle tensor. Each row represents most significant HOGC subtenors of all individual tensors at same significant level. For example 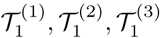 and 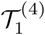 are first and most significant sub-tensors in individual tensor 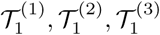 and 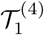 respectively in the overall tensor of cell expression, cell cycle, development and biosynthesis tensor respectively. Each path shows the common genes with higher order correlations among these sub-tensors. Each HOGC path is illustrated by different color. The left side box shows the path color for each HOGC. For example purple path passes through 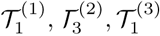 and 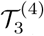 shows correlation among genes *HTB*1, *HTA*1, and *DSE*1 which are common in first sub-tensors of 𝒯^(1)^ and 𝒯^(3)^ and third significant sub-tensors of 𝒯^(2)^ and 𝒯^(4)^.

**Figure 7:**
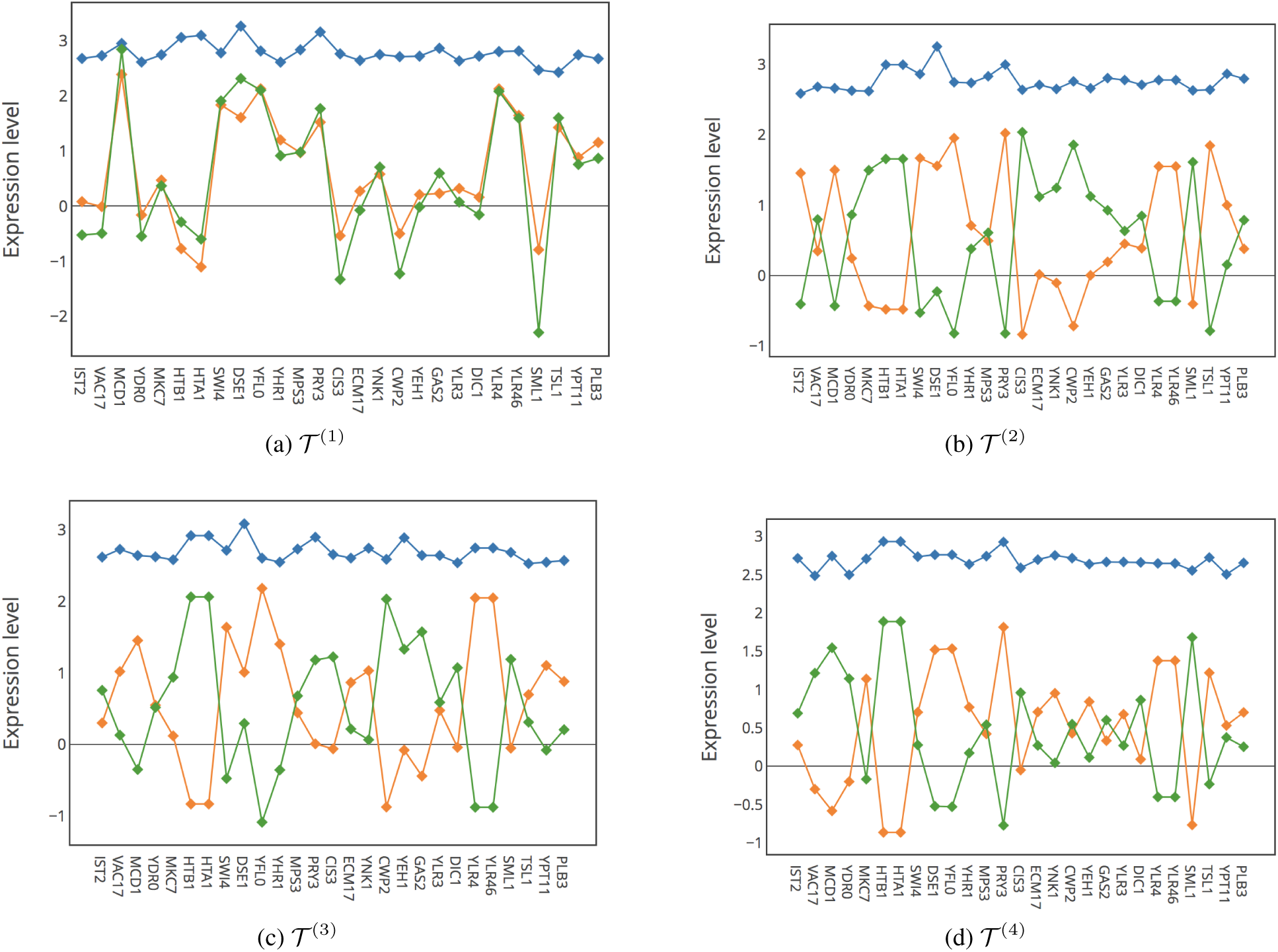
Expression patterns across gene-components in all individual tensors of 𝒯^(1)^, 𝒯^(3)^, 𝒯^(3)^and 𝒯^(4)^. In each subfigure blue line, green line, and orange one represent expression level of genes in first, second, and third most significant HOGC sub-tensor respectively in that individual tensor.

**Figure 8:**
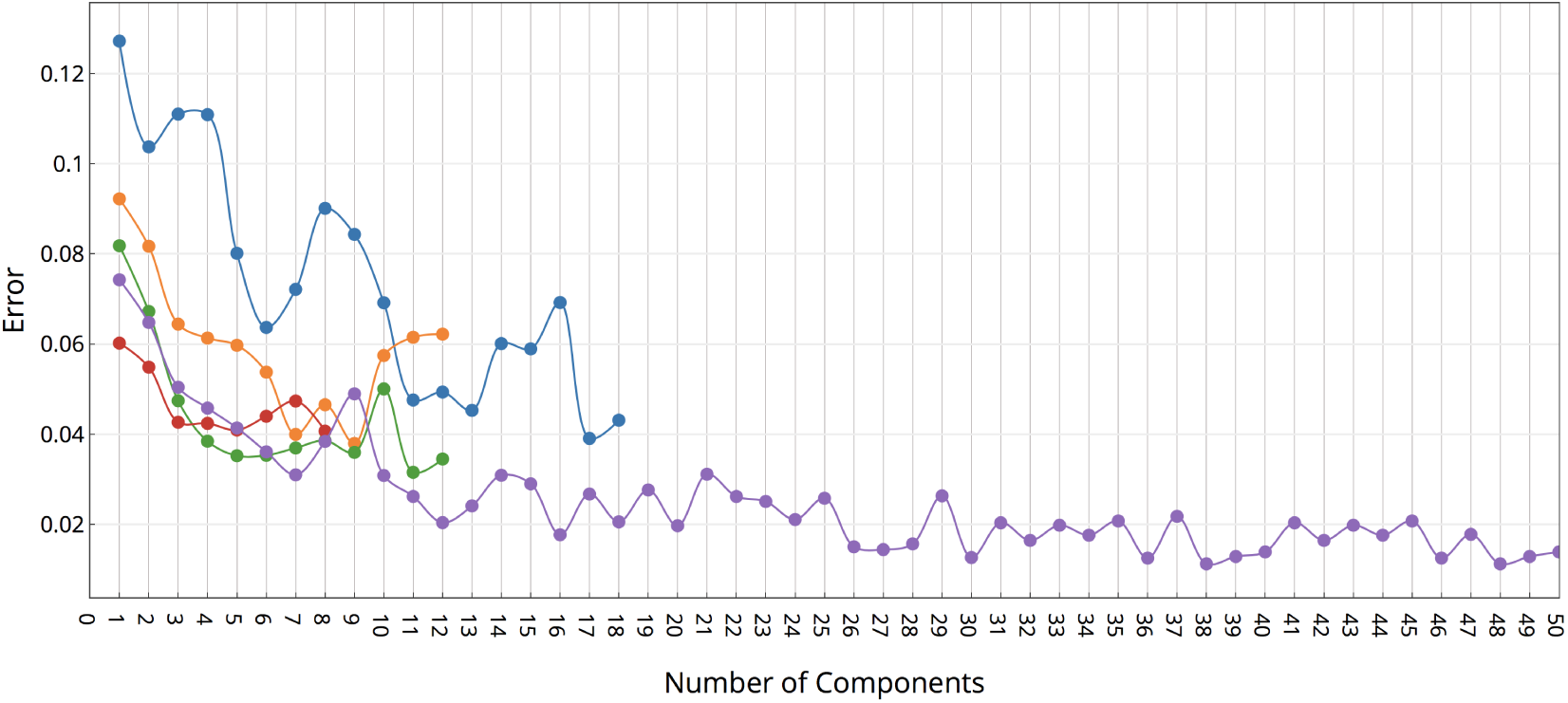
Line-joined display of spectral error of decomposition after each iteration to find a gene-components in tensors 𝒯^(1)^ (blue), 𝒯^(2)^ (orange), 𝒯^(3)^ (green), and 𝒯^(4)^ (red) and 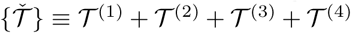 (purple). The error is the spectral norm of the deflated tensor at each step.

**Table 3:**
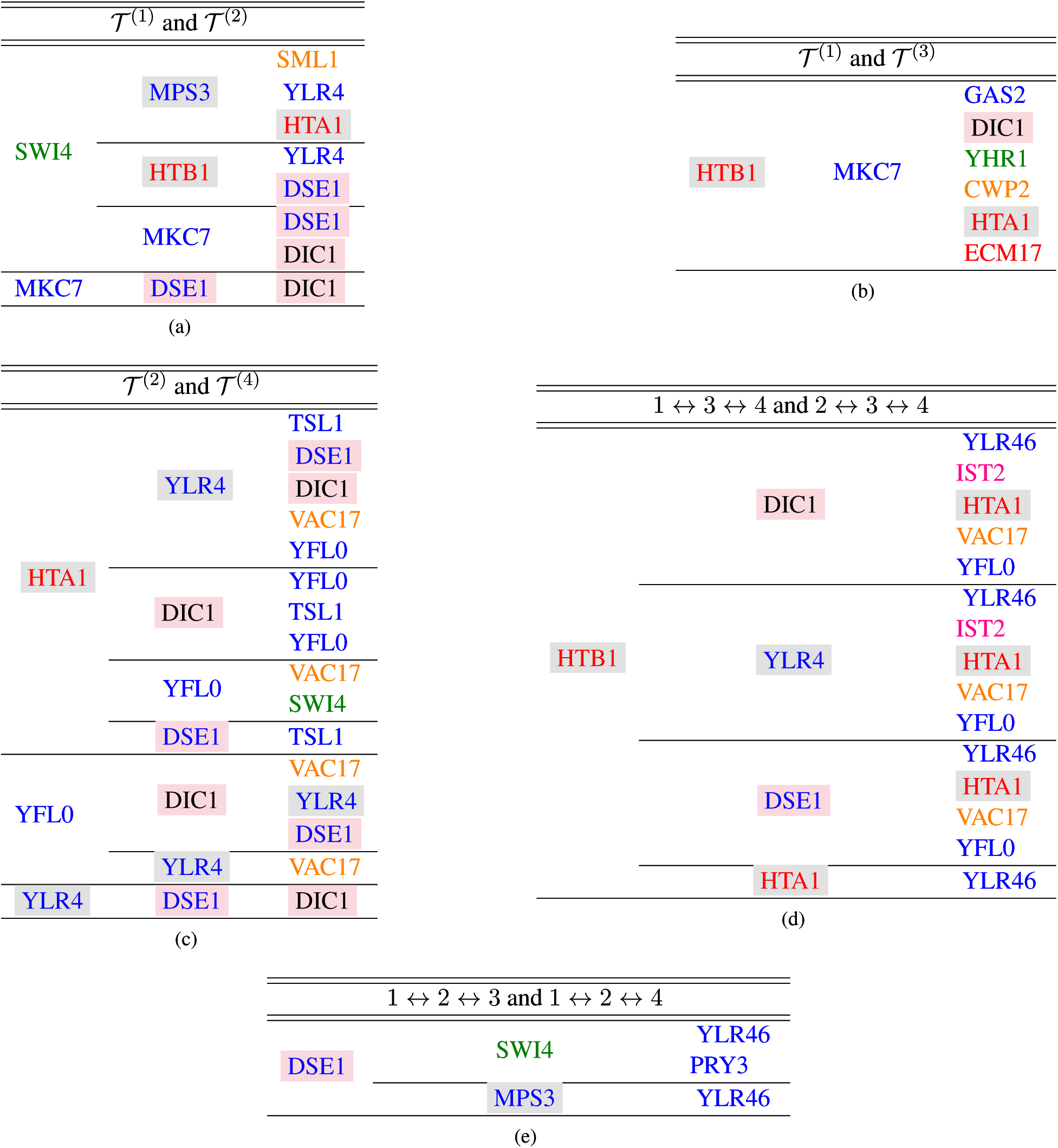
Boolean AND intersections of discretized significant sub-tensors in 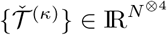 of the series of 4 individual tensors {𝒯^(1)^,𝒯^(2)^,𝒯^(3)^, 𝒯^(4)^} and their tripling in the subset of 120 higher correlations in largest amplitude among all traditionally-classified cell cycle genes of highlighted series of inter-pathway relations of the cellular system of yeast. Color-coded according to their cell-cycle classifications, *M/G*_1_(green), *G*_1_(blue), *S*(red), *S/G*_2_(orange), and *G*_2_*/M* (pink), and separately according to their pheromone-response classifications, we highlight the up-regulated purple and down-regulated gray. Yet, the annotation of *DIC*1 in cell-cycle is not reported. Our model finds the total number of 15 extracted higher-correlations of *DIC*1 with other genes among 120 higher correlations in sub-tensors. In 14 of these HOGC, one of the genes is at least from *G*_1_ stage, and the other one shows HOGC of f *HTB*1, *HTA*1, and *DIC*1 where *HTB*1 and *HTA*1 encode *S*. Therefore, our analyses predicts that *DIC*1 encodes *G*_1_under experimental conditions of Spellman et al. and Roberts et. al. [20, 21].

**Figure 9:**
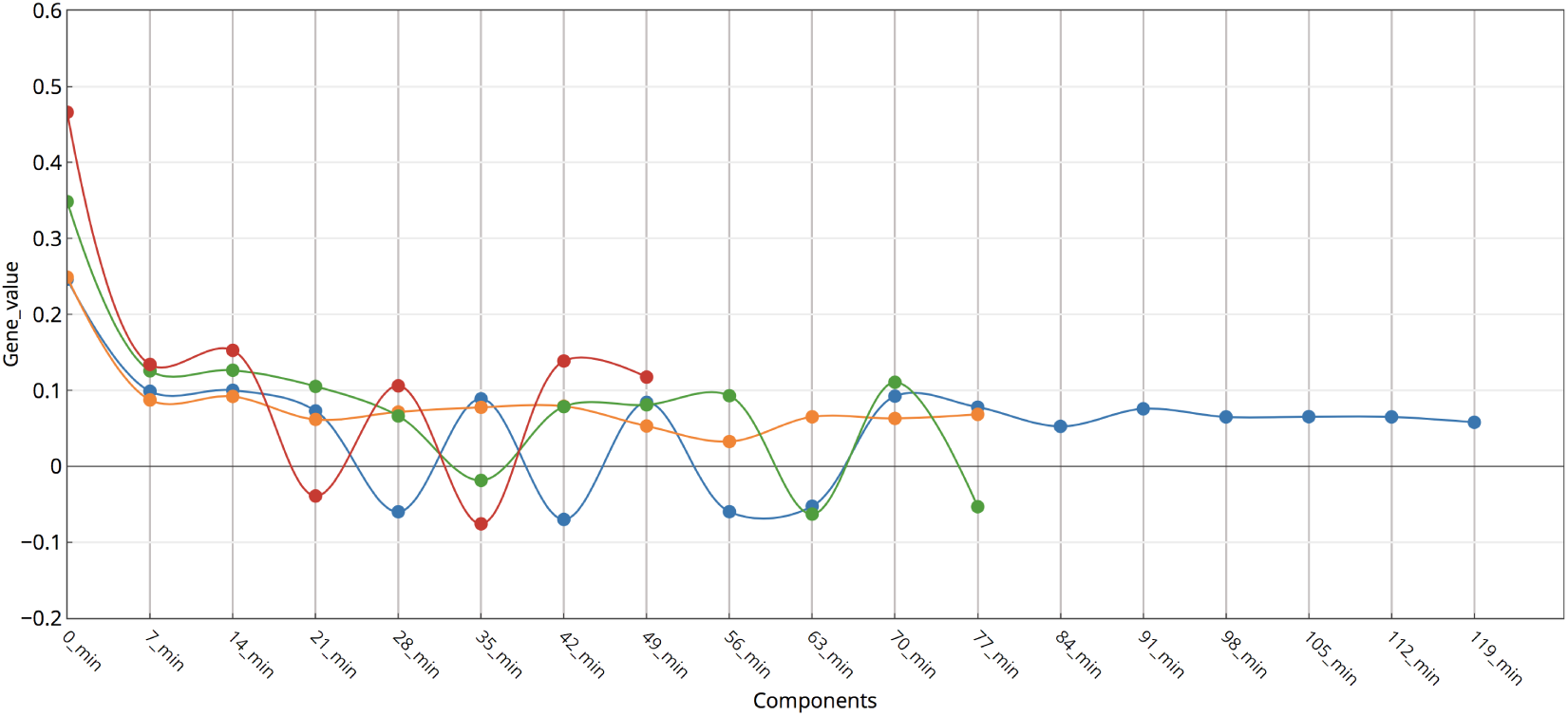
Line-joined display of gene-values for tensors 𝒯^(1)^ (blue), 𝒯^(2)^ (orange), 𝒯^(3)^ (green), and 𝒯^(4)^ (red). The *m*^*th*^ gene-value of corresponding gene-component 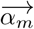 in tensor 𝒯^(*κ*)^ is 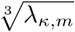 Note that for better graphical representation of it we scaled down the gene-values to third root of it which doesn’t effect on the interpretation. Each tensor is decomposed to the number of designated signal’s samples rank one sub-tensors. We find these sub-tensors and corresponding gene-value iteratively. These gene-values for each HOGC sub-tensor is computed in line 8 in Algorithm 1.

**Figure 6:**
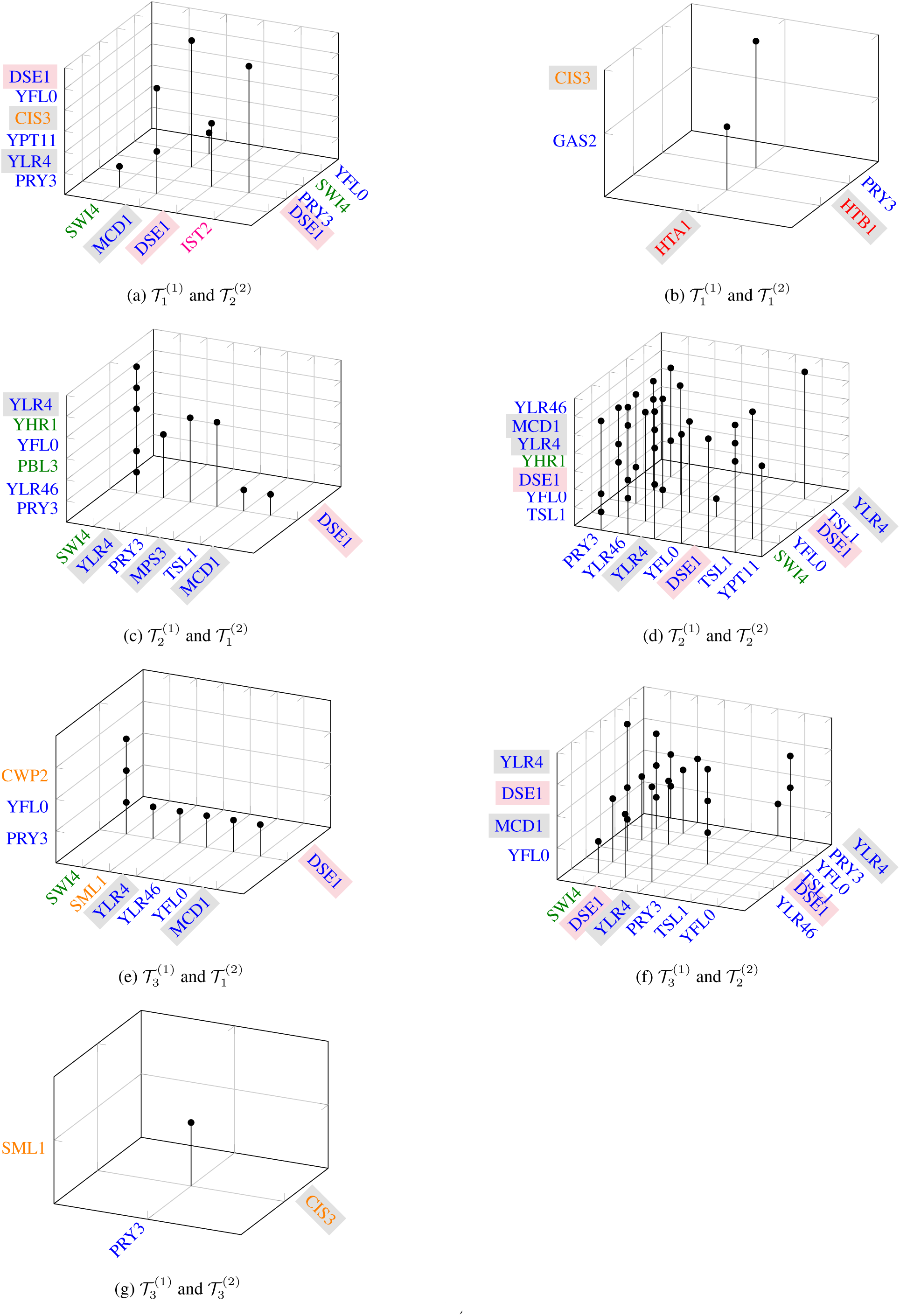
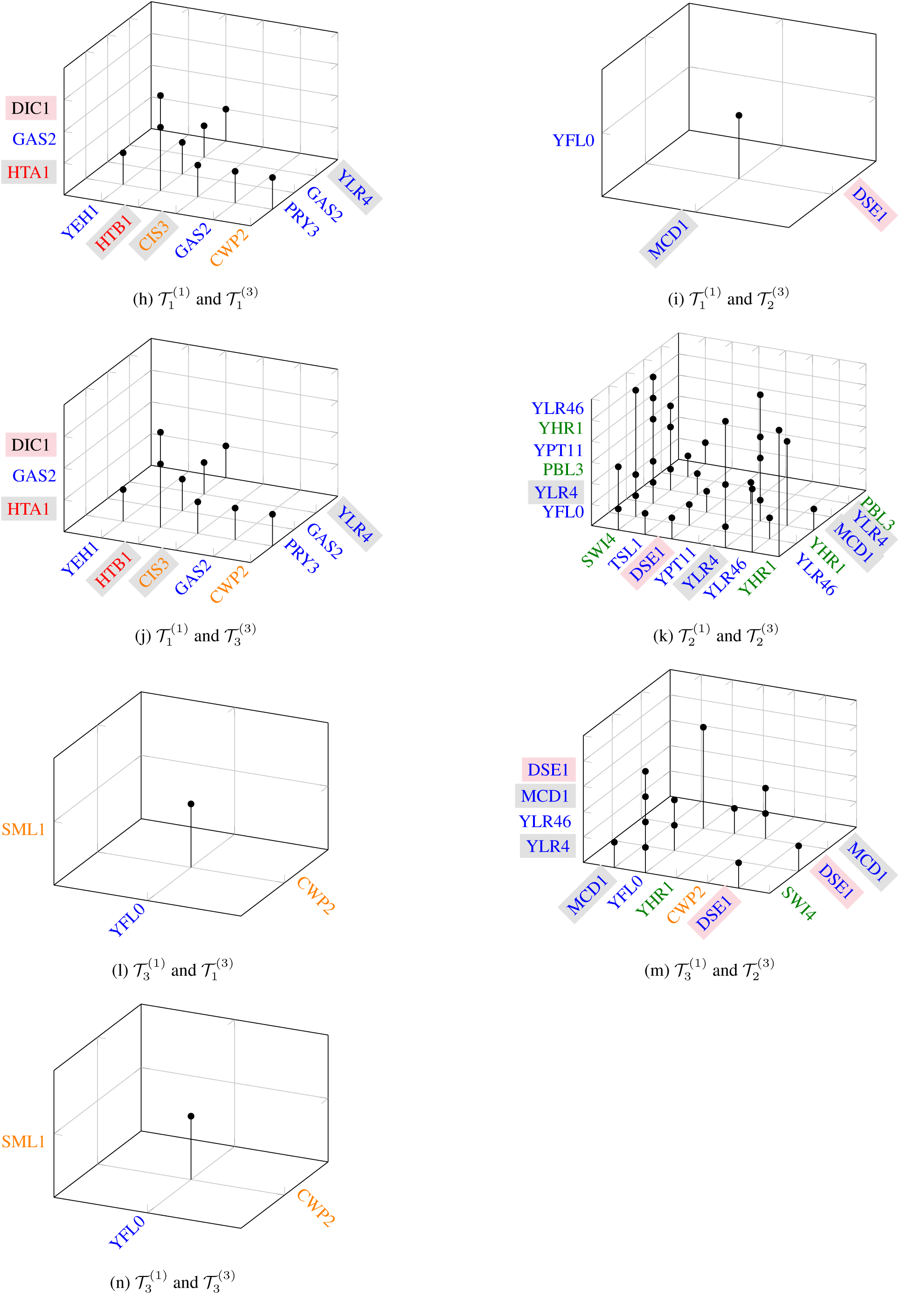
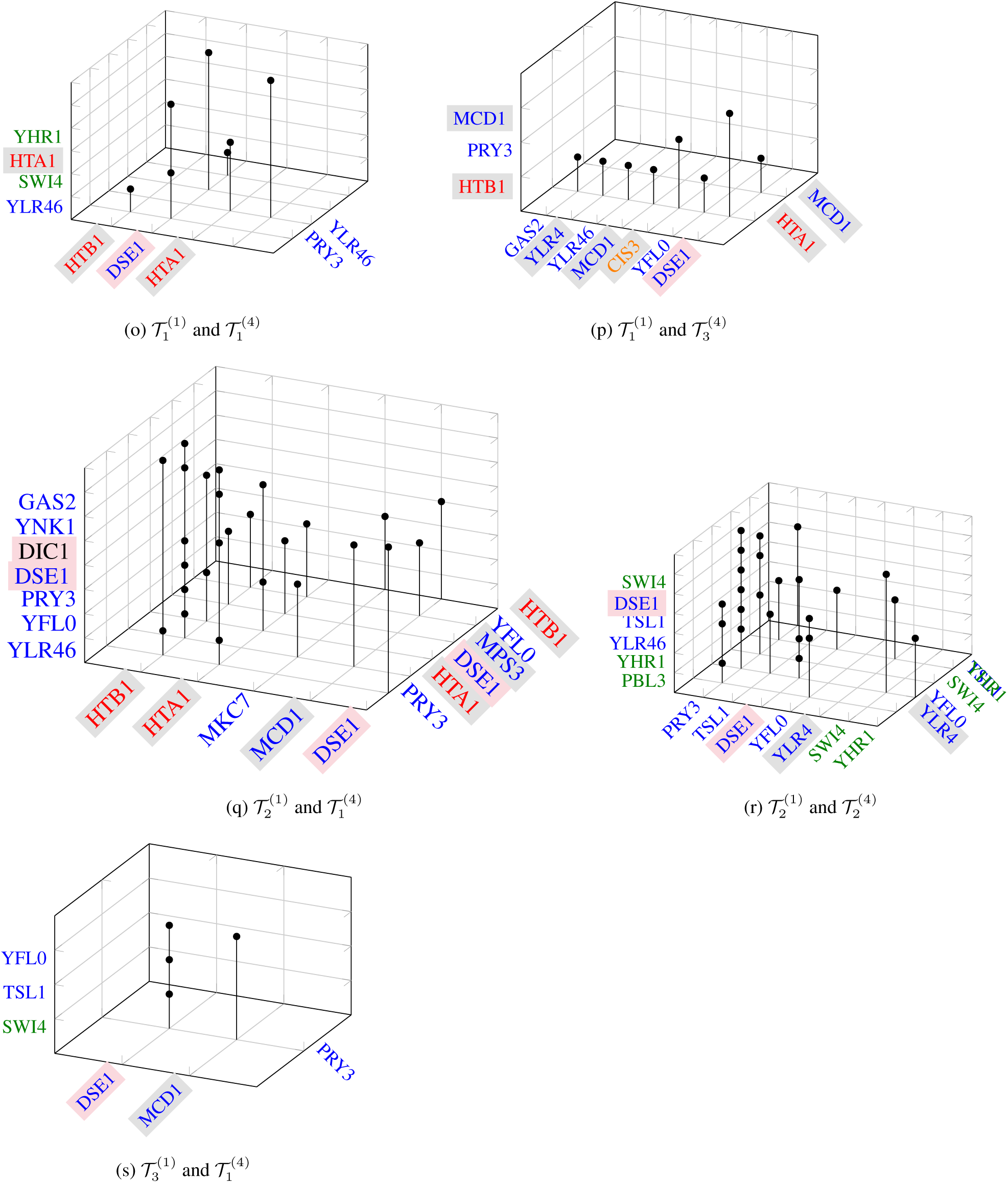
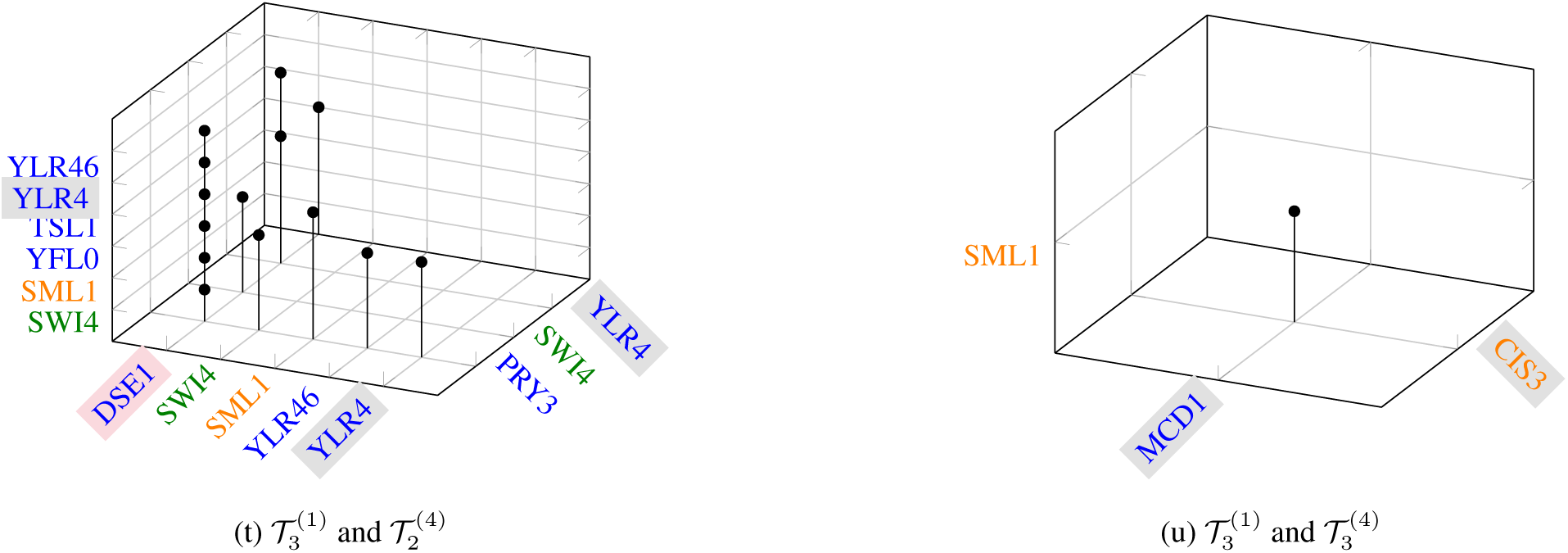
Intersection of significant HOGC sub-tensors of individual tensors in the subset of 120 relations among largest amplitude among all traditionally classified cell cycle genes of higher correlations inter-pathway dependent relations of the cellular system of yeast. Color-coded according to their cell-cycle classifications, *M/G*_1_(green), *G*_1_(blue), *S*(red), *S/G*_2_(orange), and *G*_2_*/M* (pink), and separately according to their pheromone-response classifications, we highlight the up-regulated purple and down-regulated gray. (g), (n), and (u) represents intersection of the third significant HOGC sub-tensors where two genes of three genes are with *S/G*2 and one gene is with *G*1 cell-cycle classification in all higher order signaling. The intersection uncovers higher order correlations among cell wall genes. For example *CIS*3 or *CWP* 2. Higher order correlations in third significant individual tensors would follow the same pathway-dependence. For example in all HOGC such as both *SML*1, *PRY* 3, and *CIS*3 and *SML*1, *MCD*1, and *CIS*3, we have at least one gene with down-regulated response to pheromone with cell-cycle stage *S/G*2. Therefor, this analysis suggests that *CWP* 2 in HOGC *SML*1, *Y FL*0, and *CWP* 2 which is not reported to be regulated by pheromone is down-regulated in response to pheromone. (d), (k), and (r) intersection of all second significant HOGC sub-tensors in individual tensors of 𝒯highlights higher correlation among genes with *G*_1_ and *M/G*_1_.

## Footnotes

1 *Y DR*0, *Y FL*0, *Y HR*1, *Y LR*3, *Y LR*4, and*Y LR*46 are short form of YORFs *Y DR*089*W, Y FL*064*C, Y HR*126*C, Y LR*345*W, Y LR*4262*W,* and *Y LR*4264*W* since the genes name in our data were unknown.

